# Improving peripheral reading with non-invasive transcranial electrical stimulation of early visual areas

**DOI:** 10.1101/2023.12.08.570898

**Authors:** Andrew E. Silva, Melanie A. Mungalsingh, Louise Raudzus, Benjamin Thompson

## Abstract

We investigated the ability of two non-invasive transcranial electrical stimulation (tES) protocols targeting early visual areas to improve peripheral visual word recognition in separate within-subject, double-blind, sham-controlled experiments with normal observers. English sentences were presented 10 degrees below fixation using a rapid serial visual presentation (RSVP). Transcranial random noise stimulation (tRNS) applied bilaterally on either size of the occipital pole (Oz) elicited a significant performance benefit, but transcranial direct current stimulation (tDCS) applied to Oz did not. These results highlight important factors that may influence the effectiveness of non-invasive brain stimulation methods for enhancing peripheral processing of text, such as stimulation type. While more work is necessary to better understand the relationship between tES and peripheral reading, the present results contribute to the growing body of work implicating tES as a potential tool for enhancing peripheral visual processing.

## Introduction

In peripheral vision, distinguishing an object within a cluster is more difficult than distinguishing an object presented alone, an effect called visual crowding (Bouma, 1970). Visual crowding is particularly challenging for people with central vision loss due to conditions such as macular degeneration who rely on peripheral vision to recognize objects, faces, and written text (Wallace et al., 2017). Many strategies for reducing visual crowding involve the introduction of distinguishing features to the individual objects composing the crowded visual scene. For example, crowding is reduced when the target and flanking objects have different colors or contrast polarities and when their spatial separation is increased (Kennedy & Whitaker, 2010; Levi, 2011; Scolari et al., 2007; Whitney & Levi, 2011). While between-letter and between-word crowding may negatively impact peripheral reading ability, color, contrast and spacing strategies for crowding reduction are ineffective at improving peripheral reading (Chung, 2002; Chung & Mansfield, 2009). Other studies have examined whether temporally separating the presentation of letters within a trigram or word improves overall letter or word recognition, again with limited success when using fast speeds relevant to reading (Haberthy & Yu, 2016; Silva et al., 2023).

Applying non-invasive transcranial electrical stimulation (tES) to early visual areas can reduce the strength of visual crowding without changing the visual characteristics of the stimulus, enabling better processing of cluttered visual scenes (Bello et al., 2023). The effect of tES on visual crowding has been examined in participants with normal vision (Chen et al., 2021; Contemori et al., 2019; Raveendran et al., 2020) as well as in patients with visual dysfunction, including macular degeneration (Raveendran et al., 2021), and amblyopia (Donkor et al., 2021). While visual cortex tES has demonstrated some promise for enhancing peripheral reading within a single session (Silva et al., 2022), the effect of different kinds of tES is unclear. One common tES protocol, known as transcranial direct current stimulation (tDCS), involves affixing electrodes to the head and passing a weak electrical current (1-2 mA) through the skull, stimulating the underlying neurons. The electrical current enters the body through the anode and exits through the cathode. TDCS reduces the concentration of the inhibitory neurotransmitter GABA beneath the stimulating anode, increasing cortical excitability within the local area (Stagg et al., 2009). In contrast, the area beneath the stimulating cathode exhibits reduced cortical excitability. Consequently, studies requiring increased cortical excitability will often place the cathode on a theoretically irrelevant part of the body, such as the cheek (Reinhart et al., 2016).

Although anodal tDCS applied to the visual cortex reduces crowding (Chen et al., 2021; Raveendran et al., 2020), other work has suggested that transcranial random noise stimulation (tRNS) may be a more effective modulator of cortical excitability (Fertonani et al., 2011; Inukai et al., 2016; Moliadze et al., 2014; van der Groen et al., 2022). During tRNS, the current direction rapidly alternates such that the anode and cathode identities become arbitrary, and both electrodes exert equivalent neuromodulatory effects. The alternation frequency typically ranges between 0.1 Hz and 640 Hz to coincide with physiologically measured brain oscillations (Terney et al., 2008). Protocols exclusively employing higher oscillation frequencies (100-640 Hz), known as high frequency tRNS (hf-tRNS), may influence cortical excitability more than protocols employing lower frequencies (Fertonani et al., 2011; Terney et al., 2008), though the optimal frequency range is not yet known (Moret et al., 2019).

The method of action for tRNS is currently unclear. One commonly proposed mechanism for the immediate effect of online tRNS is called stochastic resonance, whereby weak subthreshold signals are boosted by a small amount of noise to reach the threshold for detection (Moss et al., 2004). Some previous work has supported this hypothesis, demonstrating increased detection for near-threshold visual targets when the visual cortex was stimulated with tRNS at a particular strength and worse performance when other tRNS strengths were used (Groen & Wenderoth, 2016; Pavan et al., 2019), though this effect has not always been replicated across sensory modalities (Rufener et al., 2020).

One recent study examined the effect of visual cortex tDCS on reading in participants with central vision loss due to macular degeneration (Silva et al., 2022). To limit the impact of eye movements, sentences were presented on a computer screen using rapid serial visual presentation (RSVP) in which individual words were presented sequentially at specific presentation speeds and text sizes. A differential effect of tDCS was found whereby English readers exhibited a modest improvement after active tDCS but readers of Chinese exhibited no improvement (Silva et al., 2023).

Given that hf-tRNS (1) potentially modulates cortical processing more strongly than tDCS and (2) allows both electrodes to contribute to the desired neuromodulatory effect, it may be particularly effective for enhancing peripheral reading. In the current study, we tested this hypothesis in participants with normal vision across two double-blind, placebo-controlled, within-subject experiments examining the effect of hf-tRNS (Experiment 1) and tDCS (Experiment 2) on peripheral RSVP reading.

## Materials and Methods

### Participants

Eleven participants were recruited in Experiment 1 (6 females and 1 nonbinary, age mean: 23 years, age SD: 4 years). Ten different participants were recruited in Experiment 2 (6 females, age mean: 21, age SD: 5 years). All participants had normal or corrected-to-normal vision and wore their regular spectacles or contact lenses during every session. In addition, all participants were required to exhibit no contraindications for brain stimulation. All research procedures received ethics clearance from the University of Waterloo Office of Research Ethics and adhered to the tenets of the Declaration of Helsinki. All participants provided written informed consent.

### Stimulus and Apparatus

#### Rapid Serial Visual Presentation (RSVP) Task

The RSVP test was custom written using the Psychopy library in Python (Peirce et al., 2019). The stimulus was presented on an ASUS VG279 27” monitor at a pixel resolution of 1920 × 1080 and a refresh rate of 60 Hz. The viewing distance was 65 cm. Participants performed the reading task binocularly. An RSVP trial began by displaying a black fixation cross against a bright background (approximately 150 cd/m^2^). Participants were instructed to fixate on the cross for the full duration of each trial. On every trial, a randomly selected sentence was presented one word at a time 10° degrees below fixation in black Times New Roman font at a particular text size and presentation speed (Harland et al., 1998). In addition, a mask of “xxxxxxxxx” appeared before and after the sentence with identical speed and size (See Figure 1). Participants read aloud as many words as possible. The response was self-timed. Corrections were permitted but participants were required to commit to one answer per presented word. The Gazepoint GP3 eye tracker (www.gazept.com) was used to track fixation. Any trials with eye movements impinging more than 2° toward the presented word were repeated using a new sentence.

**Figure 1:**
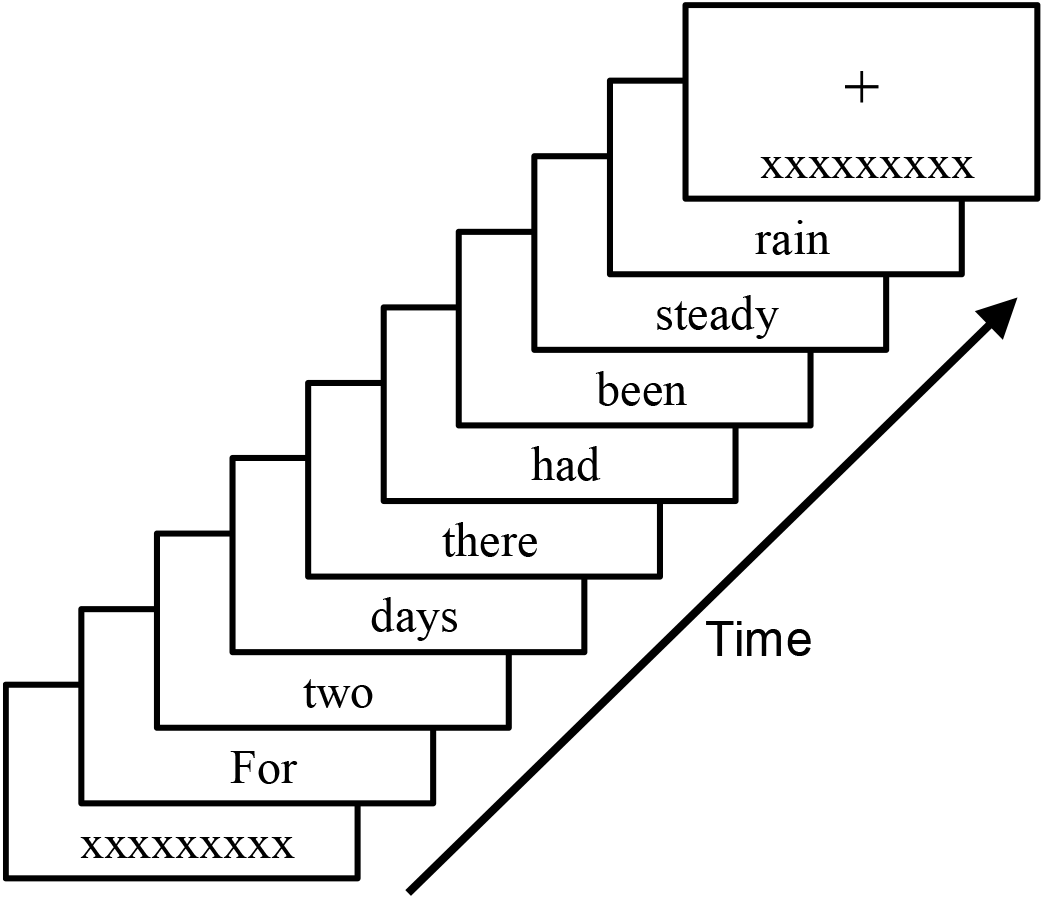
Example stimulus of one RSVP sentence. Participants were instructed to fixate on the central cross while a sentence was presented one word at a time 10° below fixation, bounded by “xxxxxxxxx”. The task was to read the sentence aloud, and performance accuracy was calculated as the total proportion of words read correctly across all presented sentences exhibiting the same presentation speed and text size.

All sentences were selected randomly from a pool of 2,627 sentences extracted from nine classic novels written in English. The total length of all the sentences used ranged between 40- and 80-characters including spaces (Chung et al., 1998). No individual sentence was seen more than once by any participant.

#### Transcranial electrical stimulation

Participants underwent one active tES session and one placebo tES session. The order of sessions was randomly selected and counterbalanced. Neither the researcher nor the participant was informed of the order of sessions. The active session was either tDCS (Experiment 1: 2 mA, 20 min, 30 sec ramp up/down) or hf-tRNS (Experiment 2: 2 mA peak to peak, 20 minutes, 30 seconds ramp up/down). The placebo session consisted only of 30 second ramp up and ramp down periods. The current was delivered with a neuroConn DC Stimulator Plus (neurocaregroup.com) using two 5 cm x 5 cm rubber electrodes placed inside saline-soaked sponges that were attached to the head with elastic bands.

During the Experiment 1 hf-tRNS sessions, both electrodes were placed on either side of Oz lengthwise, with the electrode cables exiting outward. The electrodes were placed 2-3 finger widths apart, and care was taken to maintain a dry section of scalp between electrodes. During the Experiment 2 tDCS sessions, the anode was placed over Oz using the International 10-20 EEG system. The cathode was placed over a randomly selected cheek which remained consistent across both tDCS sessions.

### Experimental Procedure

Participants performed three experimental sessions. In the first session, a threshold reading speed and size was found without tES. In the second and third sessions, the main experimental task was performed using the text size and speed found during the first session.

#### Session 1: Thresholding and parameter selection

The purpose of the thresholding session was to find an appropriate text size and speed for use during the tES sessions that would elicit 55% performance, representing a challenging threshold far from ceiling and floor performance. During thresholding, participants performed the RSVP reading task with five initial text sizes, each with five initial presentation speeds (Table 1). These values were selected from prior pilot testing. Four trials per condition were run, totalling 75 initial trials. Performance accuracy was calculated as the proportion of words identified correctly across all trials exhibiting the same combination of text size and presentation speed. The text sizes were block randomized, and the presentation speeds were randomly interleaved within blocks.

**Table 1:** Initial text sizes and presentation speeds presented to each participant during Session 1 thresholding.

|  | Print Size (logMAR) |  |  |  |  |
| --- | --- | --- | --- | --- | --- |
|  | 0.92 | 1.08 | 1.23 | 1.36 | 1.43 |
| Presentation | 83 | 50 | 50 | 50 | 50 |
| Speeds | 150 | 100 | 100 | 100 | 100 |
| (ms) | 300 | 250 | 200 | 150 | 133 |
|  | 700 | 450 | 400 | 250 | 233 |
|  | 2500 | 1000 | 1000 | 800 | 700 |

logMAR = logarithm of the minimum angle of resolution, ms = millisecond

The presentation speeds were selected to elicit performance spanning 20%-80% accuracy.

Directly after completing the initial 75 trials, the data from each text size were fit to separate cumulative gaussian psychometric functions describing performance with changing presentation speed (See Figure 2A-E for representative psychometric fits). If performance on any text size failed to span the required 20%-80% performance accuracy, then additional trials were run with new reading speeds to span the full psychometric function.

**Figure 2:**
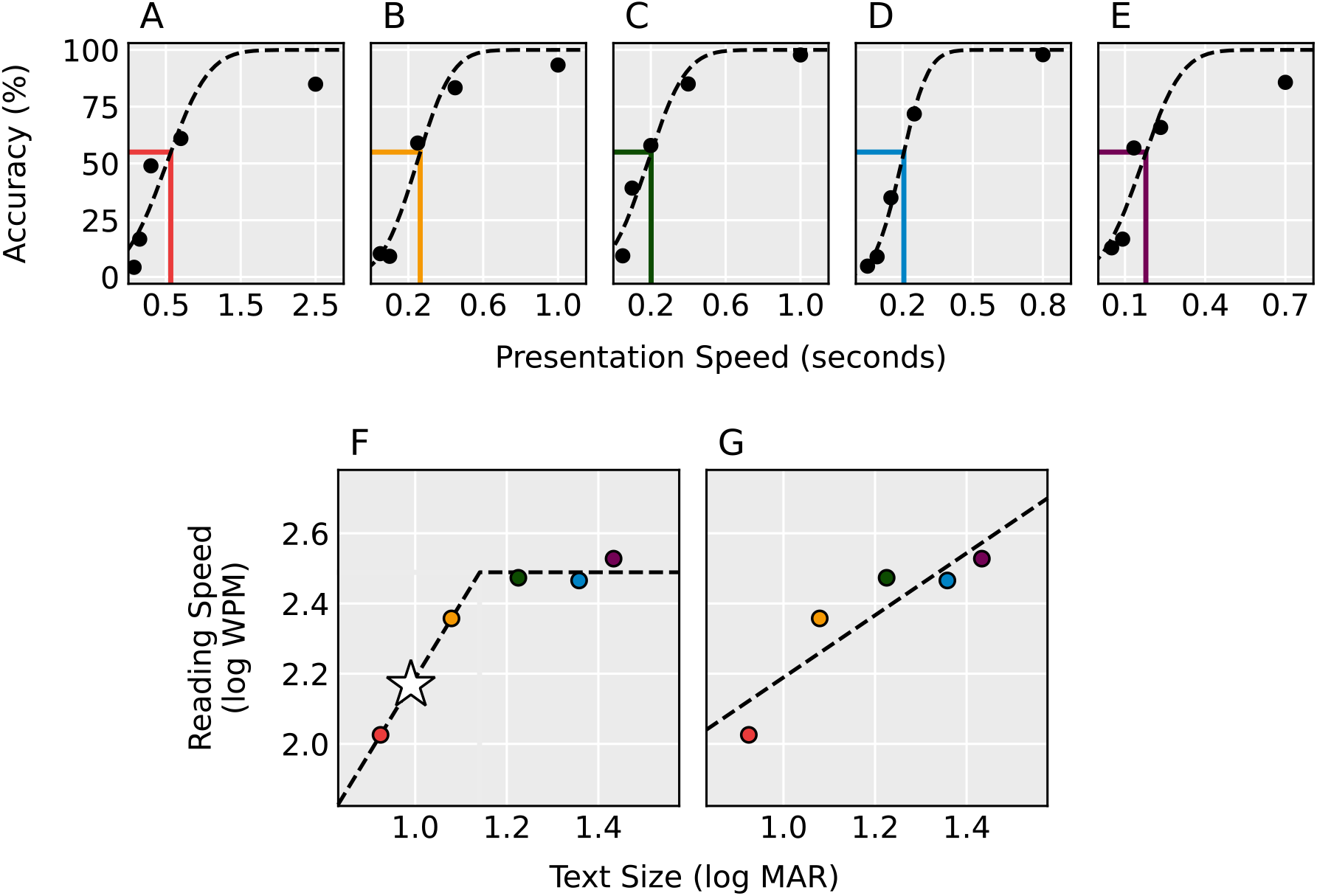
Baseline thresholding procedure for one example participant. Participants performed the RSVP task with 5 different text sizes (logMAR, A: 0.92, B: 1.08, C: 1.23, D: 1.36, E: 1.43) and a cumulative gaussian psychometric function was fit to the data from each size. The presentation speeds eliciting 55% accuracy for each text size were fit to a continuous piecewise function with one increasing segment and one flat segment (F) and a straight line (G). Additional print sizes were run until the piecewise function fit the data better than the straight line. The text size and reading speed used during the stimulation sessions was selected by taking the point along the increasing segment corresponding to 0.15 logMAR below the intersection of both segments (star).

In addition, tested print sizes (logMAR) and their associated presentation speeds (log_10_ words per minute) eliciting 55% accuracy were fit to a standard linear function and a continuous bilinear piecewise function directly after completing the initial 75 trials (Chung et al., 1998). The piecewise funct on conta ned one segment that r ses w th ncreas ng pr nt s ze up to a max mum “cr t cal pr nt s ze” (CPS) and one hor zontal segment eg nn ng at the cr t cal pr nt s ze and extend ng to all larger print sizes. The CPS and associated presentation speed demarcates the smallest text with which the participant can achieve their maximum reading speed when targeting a specific accuracy (55% in the current study). The sum of the squared residuals of both models were calculated to verify a reliable CPS. If the linear fit explained equal or greater amounts of the variance compared to the piecewise fit, then an additional print size was run to better capture the CPS. This procedure occurred iteratively until the piecewise fit explained more of the variance. See Figure 2F and 2G for representative linear and piecewise fits.

Session 1 was completed after the CPS was found. The print size and presentation speed along the estimated piecewise function 0.15 logMAR below the CPS were selected for use during the part c pant’s tES sessions. The selected parameters fall along the sloped portion of the piecewise function and are therefore theoretically sensitive to improvements in both visual acuity and reading speed.

#### Sessions 2 & 3: tES experiments

For any given participant across both experiments, Sessions 2 & 3 were identical, except that one session provided active tES and the other session provided placebo tES in a randomly selected order. The experimenters and participants were both blinded to the order of stimulation. Sessions 2 & 3 were separated by at least 2 days and no more than 7 days. On each session, four separate RSVP reading tests were administered. Each test comprised 30 sentences, and performance accuracy was calculated as the total proportion of words identified correctly. The presentation speed and text size were determ ned Sess on 1’s parameter select on procedure and rema ned constant throughout both stimulation sessions. All other RSVP procedures were identical to Session 1.

Prior to tES, participants performed an RSVP pre-test. After the pre-test was concluded, either active or placebo tES was applied (Experiment 1: hf-tRNS applied bilaterally to early visual areas, Experiment 2: tDCS applied with the anode over Oz and the cathode over a randomly selected cheek). A second RSVP test was administered during tES stimulation (“dur ng”). A third test was administered 5 minutes after the st mulat on completed (“post5”). The f nal test was adm n stered 30 m nutes after the st mulat on completed (“post30”).

## Results

The effects of active and placebo tES were calculated by subtracting the pre-test accuracy from each of the post-test accuracies. Separate 2 (stimulation type: Active and Placebo) × 3 (time: During, Post5, Post30) repeated-measures ANOVAs were run on the normalized data from Experiment 1 and Experiment 2. All analyses were carried out using jamovi v2.3(www.jamovi.org).

### Experiment 1 - tRNS

One outlier was found using a 1.5 × interquartile range threshold that exhibited an exceedingly large difference between the effects of active hf-tRNS and placebo. This anomalous result was driven by poor performance during the placebo post-tests relative to pre-test (See Table 2). Therefore, this participant was omitted from the statistical analysis.

**Table 2:** Individual Subject Data, Experiment 1 (hf-tRNS).

| Subject ID | Active Session Accuracy (%) |  |  |  | Placebo Session Accuracy (%) |  |  |  |
| --- | --- | --- | --- | --- | --- | --- | --- | --- |
|  | Pre-Test | During | Post5 | Post30 | Pre-Test | During | Post5 | Post30 |
| 1 | 89 | 96 | 94 | 96 | 92 | 97 | 91 | 93 |
| 2 | 82 | 82 | 77 | 82 | 80 | 77 | 80 | 79 |
| 3 | 81 | 83 | 87 | 86 | 74 | 70 | 72 | 81 |
| 4 | 62 | 66 | 67 | 68 | 58 | 60 | 59 | 60 |
| 5 | 80 | 84 | 83 | 89 | 80 | 79 | 83 | 82 |
| 6 | 54 | 60 | 51 | 51 | 57 | 55 | 52 | 47 |
| 7 | 84 | 89 | 85 | 84 | 79 | 72 | 83 | 82 |
| 8 | 88 | 90 | 81 | 88 | 82 | 76 | 79 | 85 |
| 9 | 52 | 69 | 77 | 76 | 56 | 66 | 66 | 70 |
| 10 | 84 | 91 | 95 | 90 | 92 | 94 | 91 | 96 |
| ^11 | 64 | 63 | 67 | 74 | 76 | 41 | 47 | 55 |

hf-tRNS = High frequency transcranial random noise stimulation.
% = Percent Correct.
During = Reading test performed while stimulation is administered.
Post5 = Reading test performed 5 minutes after stimulation completed.
Post30 = Reading test performed 30 minutes after stimulation completed.
^ Outlier data as defined by a 1.5 × IQR threshold on the different between the pre and post tests.

Active stimulation significantly improved performance relative to placebo stimulation, *F*(1,9) = 19.6, *p* = 0.002 (active mean: 4.8%, active SEM: 2.1, placebo mean: 0.5%, placebo SEM: 1.4).

There was no significant effect of time on performance and no significant interaction. Figure 3A illustrates the Experiment 1 data at pre-test and each of the three post tests, and Figure 4 illustrates the difference between the effects of active hf-tRNS and placebo.

**Figure 3:**
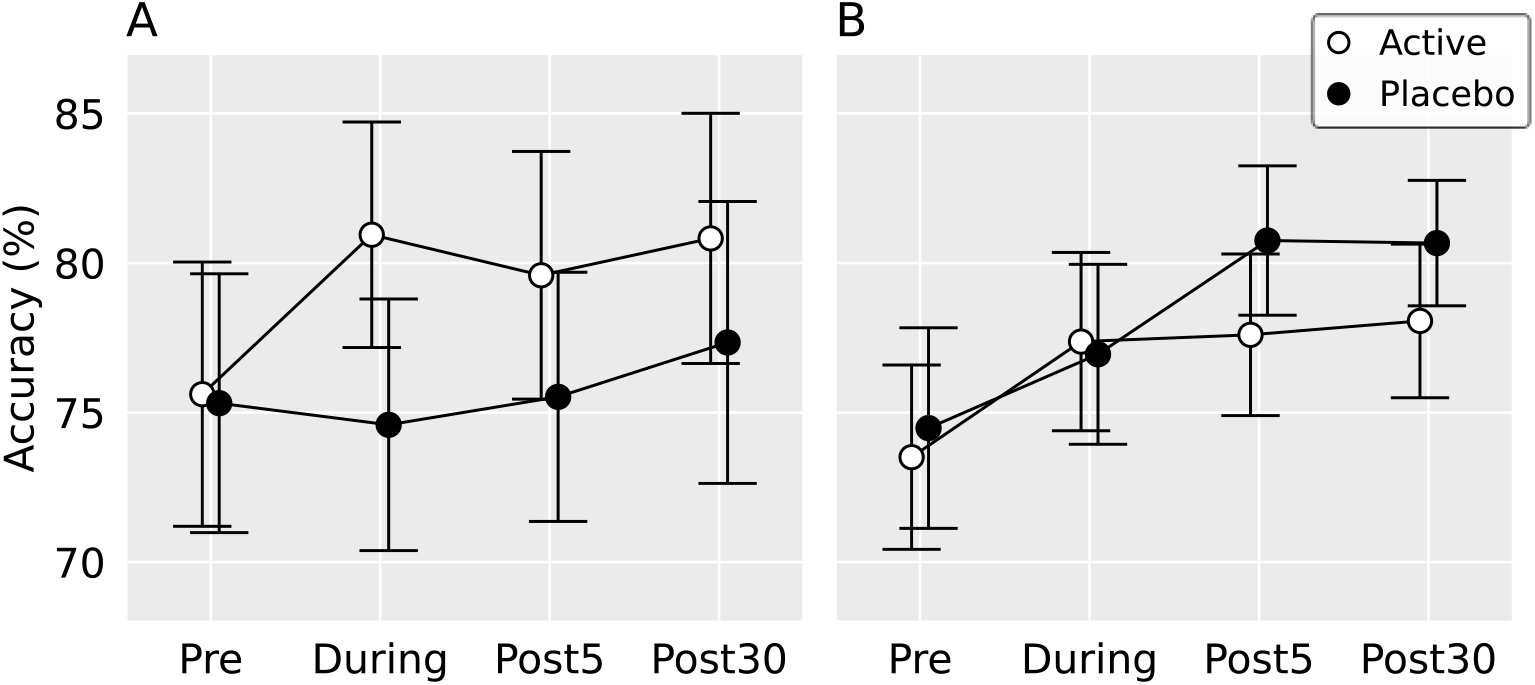
Group-averaged performance accuracies for Experiment 1 (hf-tRNS, A) and Experiment 2 (tDCS, B). The Experiment 1 outlier (Table 3) was excluded, and error bars are ±1 standard error of the mean.

**Figure 4:**
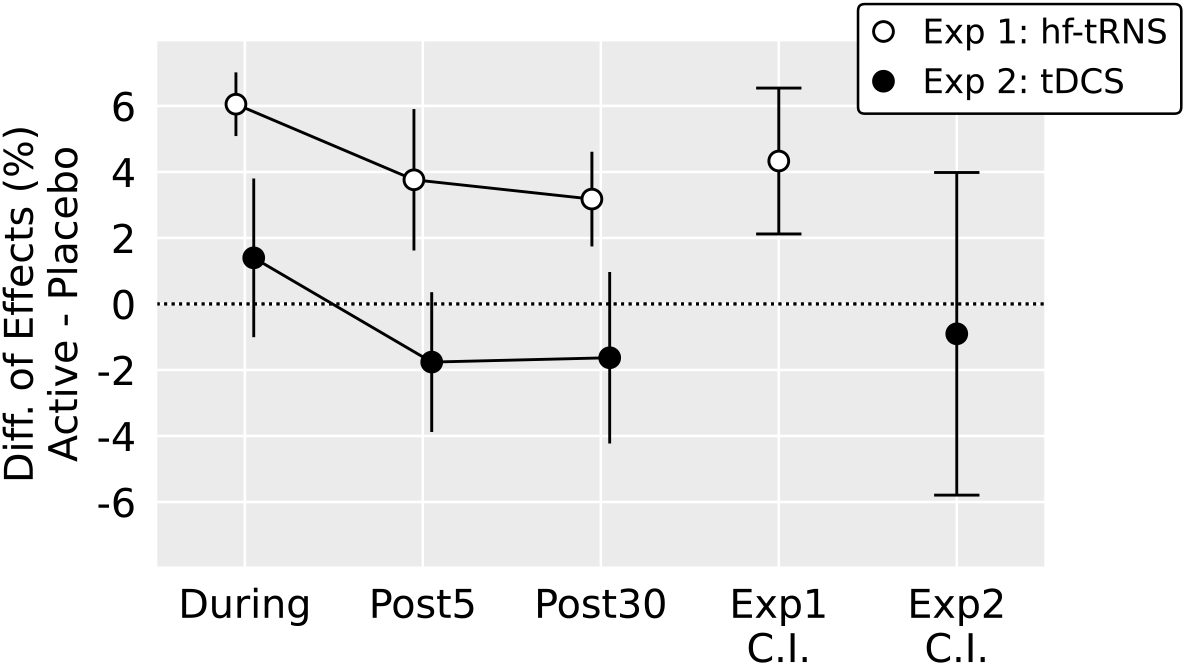
The differences between the effects of active and placebo stimulation in Experiment 1 and Experiment 2 across each baseline-normalized post-test. The right-most two points show overall 95% confidence intervals for the difference between active and placebo stimulation.

### Experiment 2 – tDCS

See Table 3 for individual subject data from Experiment 2. There was no significant main effect of stimulation type, no significant effect of time, and no significant interaction (active mean: 4.0%, active SEM: 1.5, placebo mean: 4.9%, placebo SEM: 1.4). Figures 3B and 4 illustrate the Experiment 2 data, and Table 3 presents the individual subject data.

**Table 3:** Individual Subject Data, Experiment 2 (tDCS).

| Subject ID | Active Session Accuracy (%) |  |  |  | Placebo Session Accuracy (%) |  |  |  |
| --- | --- | --- | --- | --- | --- | --- | --- | --- |
|  | Pre-Test | During | Post5 | Post30 | Pre-Test | During | Post5 | Post30 |
| 1 | 88 | 85 | 88 | 88 | 77 | 84 | 87 | 90 |
| 2 | 70 | 80 | 83 | 77 | 80 | 79 | 82 | 81 |
| 3 | 78 | 86 | 78 | 73 | 73 | 75 | 75 | 81 |
| 4 | 72 | 78 | 68 | 76 | 75 | 77 | 79 | 78 |
| 5 | 79 | 90 | 85 | 85 | 88 | 89 | 92 | 85 |
| 6 | 51 | 61 | 62 | 59 | 49 | 55 | 66 | 67 |
| 7 | 69 | 69 | 76 | 78 | 78 | 80 | 83 | 78 |
| 8 | 76 | 68 | 78 | 79 | 82 | 84 | 88 | 89 |
| 9 | 77 | 85 | / | 85 | 75 | 79 | 82 | 81 |
| 10 | 75 | 73 | 80 | 82 | 66 | 68 | 72 | 76 |

tDCS = Transcranial direct current stimulation.
% = Percent Correct.
During = Reading test performed while stimulation is administered.
Post5 = Reading test performed 5 minutes after stimulation completed.
Post30 = Reading test performed 30 minutes after stimulation completed.
/ = Data unavailable.

### Hf-tRNS vs. tDCS

As an additional analysis, we statistically compared the effects of hf-tRNS and tDCS. We calculated the overall improvement of active stimulation relative to placebo for each participant by averaging the associated During, Post5, and Post30 post-tests and subtracting the pre-test. Su ject 9’s active Post5 test was excluded from this analysis (See Table 3). We then subtracted the overall effect of placebo stimulation from the overall effect of active stimulation for each participant (See the Figure 4 confidence intervals). The resulting overall hf-tRNS and tDCS effects were compared with a two-tailed independent samples *t*-test, finding that active hf-tRNS elicited significantly better performance relative to sham than tDCS, *t*(18) = 2.27, *p* = 0.036.

## Discussion

The current study explored the effects of hf-tRNS and tDCS applied to early visual areas on peripheral reading. We found that active bilateral hf-tRNS produced a significant improvement, but anodal tDCS to Oz did not produce any benefit. Our findings suggest that hf-tRNS may functionally improve peripheral reading a result with potential relevance to people exhibiting low vision due to ocular diseases like macular degeneration. This effect may be partly driven by the ability of both tRNS stimulating electrodes to contribute to cortical neuromodulation.

The current study found no effect of tDCS. However, Silva et al. (2022) utilized a similar RSVP protocol and reported an effect with tDCS. The reason for a null tDCS effect in the current study is unclear. One of the most pronounced differences between the two studies is how visual fixation was controlled. The current study presented stimuli 10° away from fixation—a relatively large eccentricity selected because it falls beyond the macula (Strasburger et al., 2011) and has been used in previous work (Chung et al., 2004; Yu et al., 2010). In contrast, S lva et al.’s (2022) participants with macular degeneration adopted their choice of fixation eccentricity, potentially the nearest eccentric position possible. Because more eccentric retinotopic positions are located deeper within the calcarine sulcus (Wu et al., 2012), the ideal stimulation depth likely differed between the current study and Silva et al. (2022).

The effects of tDCS are impacted by cortical folding such that excitatory protocols may actually induce antagonistic excitatory and inhibitory effects on the stimulated region due to the underlying tissue structure (Terney et al., 2008). In contrast, tRNS induces strictly excitatory stimulation that is more robust to cortical folding (Terney et al., 2008), possibly allowing deeper penetration of the stimulation. TDCS’ greater dependence on a direct path to the stimulation target may have inhibited meaningful neuromodulation of the cortical region supporting visual processing at 10° eccentricity. Future investigations of the relationship between RSVP reading eccentricity and the efficacy of different tES protocols will be particularly relevant for evaluating potential clinical efficacy.

While both tDCS and tRNS are implicated in the reduction of visual crowding (Bello et al., 2023), it is also possible that the improvement due to brain stimulation found in the current study was not strictly driven by reduced visual crowding. Although peripheral RSVP reading tasks are beholden to visual crowding, they do not inherently measure visual crowding. Visual acuity is markedly worse at 10° eccentricity than at central fixation, particularly when viewing written words (Abdelnour & Kalloniatis, 2001). Consequently, the signal strength of our current stimulus is correspondingly lower and noisier than if the sentences had been centrally presented. Rather than a reduction of visual crowding, the improvement resulting from hf-tRNS may have been due to stochastic resonance (Groen & Wenderoth, 2016; Pavan et al., 2019). High frequency tRNS-induced random noise applied to the early visual areas may have boosted the weak peripheral signal, improving general visual acuity and increasing task performance independent of visual crowding. Because the current study did not explicitly measure crowding, the precise mechanism underlying the observed effect of brain stimulation remains unclear.

It should be noted that reading comprehension was not tested in this study. Rather, participants performed a visual word recognition task that is presumably less dependent on higher level cognitive processing than sentence comprehension. Because visual word recognition is dramatically impaired in the periphery (Abdelnour & Kalloniatis, 2001), it was necessary to use individualized presentation speeds and text sizes to control task difficulty across all participants. While our initial baseline thresholding procedure avoided ceiling and floor effects, a clear learning effect between the initial thresholding session and the experimental tES sessions was observed across most participants. We targeted a point below the 55% critical print size (CPS) and corresponding reading speed to enable detection of improvements in either visual acuity or reading speed during the experimental tests. Unfortunately, the elevated pre-tests (See Tables 2 and 3) indicate that we cannot e sure where the st mulus fell along each part c pant’s curve describing reading speed as a function of print size, demonstrated in Figure 2F. It would be interesting to examine whether the effect of tES can be enhanced with more optimal stimulus parameters. Nevertheless, the robust effect of hf-tRNS demonstrates that neuromodulation may be useful for enhancing visual processing in the periphery, particularly for people with low vision using the periphery to read.

Finally, our exploratory analysis found a significant difference between the effects of tDCS and hf-tRNS. However, these separate experiments were not initially designed to be directly compared and were carried out at different times. Additionally, the behavior of the sham groups differed between the two studies, again suggesting that caution should be used when directly comparing the two experiments.

## Conclusion

The current study found a robust enhancement of peripheral RSVP word recognition accuracy after hf-tRNS, but not after tDCS. Interestingly, this result runs counter to a previous study that found a significant effect with tDCS in patients with macular degeneration (Silva et al., 2022). The current study fixed eccentricity to 10°, whereas the patient study allowed participants to use their preferred retinal location for viewing. The resulting differences in stimulus eccentricity may have contributed to the pattern of results observed in this investigation. Although additional work must clarify optimal visual stimulus and neuromodulation parameters, non-invasive brain stimulation of early visual areas may be potentially useful for improving the visual processing of text presented to the periphery. In particular, tRNS appears effective for relatively far peripheral locations.

## Acknowledgements

The authors would like to thank Karen Fan, Kate Woo, and Mawj Al-Hammadi for assistance with data collection.

## References

Abdelnour, O., & Kalloniatis, M. (2001). Word Acuity Threshold as a Function of Contrast and Retinal Eccentricity. Optometry and Vision Science, 78(12), 914. 10.1097/00006324-200112000-00014

Bello, U. M., Wang, J., Park, A. S. Y., Tan, K. W. S., Cheung, B. W. S., Thompson, B., & Cheong, A. M. Y. (2023). Can visual cortex non-invasive brain stimulation improve normal visual function? A systematic review and meta-analysis. Frontiers in Neuroscience, 17. 10.3389/fnins.2023.1119200

Bouma, H. (1970). Interaction effects in parafoveal letter recognition. Nature, 226(5241), 177–178. 10.1038/226177a0

Chen, G., Zhu, Z., He, Q., & Fang, F. (2021). Offline transcranial direct current stimulation improves the ability to perceive crowded targets. Journal of Vision, 21(2), 1–1. 10.1167/JOV.21.2.1

Chung, S. T. L. (2002). The effect of letter spacing on reading speed in central and peripheral vision. Investigative Ophthalmology and Visual Science, 43(4), 1270–1276.

Chung, S. T. L., Legge, G. E., & Cheung, S. H. (2004). Letter-recognition and reading speed in peripheral vision benefit from perceptual learning. Vision Research, 44(7), 695–709. 10.1016/J.VISRES.2003.09.028

Chung, S. T. L., & Mansfield, J. S. (2009). Contrast polarity differences reduce crowding but do not benefit reading performance in peripheral vision. Vision Research, 49(23), 2782–2789. 10.1016/J.VISRES.2009.08.013

Chung, S. T. L., Mansfield, J. S., & Legge, G. E. (1998). Psychophysics of reading. XVIII. The effect of print size on reading speed in normal peripheral vision. Vision Research, 38(19), 2949–2962. 10.1016/S0042-6989(98)00072-8

Contemori, G., Trotter, Y., Cottereau, B. R., & Maniglia, M. (2019). tRNS boosts perceptual learning in peripheral vision. Neuropsychologia, 125, 129–136. 10.1016/j.neuropsychologia.2019.02.001

Donkor, R., Silva, A. E., Teske, C., Wallis-Duffy, M., Johnson, A. P., & Thompson, B. (2021). Repetitive visual cortex transcranial random noise stimulation in adults with amblyopia. Scientific Reports, 11(1), 3029. 10.1038/s41598-020-80843-8

Fertonani, A., Pirulli, C., & Miniussi, C. (2011). Random Noise Stimulation Improves Neuroplasticity in Perceptual Learning. The Journal of Neuroscience, 31(43), 15416–15423. 10.1523/JNEUROSCI.2002-11.2011

Groen, O. van der, & Wenderoth, N. (2016). Transcranial Random Noise Stimulation of Visual Cortex: Stochastic Resonance Enhances Central Mechanisms of Perception. Journal of Neuroscience, 36(19), 5289–5298. 10.1523/JNEUROSCI.4519-15.2016

Haberthy, C., & Yu, D. (2016). Effects of temporal modulation on crowding, visual span, and reading. Optometry and Vision Science, 93(6), 579–587. 10.1097/OPX.0000000000000848

Harland, S., Legge, G. E., & Luebker, A. (1998). Psychophysics of Reading. XVII. Low-Vision Performance with Four Types of Electronically Magnified Text. Optometry and Vision Science, 75(3), 183. 10.1097/00006324-199803000-00023

Inukai, Y., Saito, K., Sasaki, R., Tsuiki, S., Miyaguchi, S., Kojima, S., Masaki, M., Otsuru, N., & Onishi, H. (2016). Comparison of Three Non-Invasive Transcranial Electrical Stimulation Methods for Increasing Cortical Excitability. Frontiers in Human Neuroscience, 10(December), 1–7. 10.3389/fnhum.2016.00668

Kennedy, G. J., & Whitaker, D. (2010). The chromatic selectivity of visual crowding. Journal of Vision, 10(6), 15–15. 10.1167/10.6.15

Levi, D. M. (2011). Visual crowding. Current Biology, 21(18), R678–R679. 10.1016/j.cub.2011.07.025

Moliadze, V., Fritzsche, G., & Antal, A. (2014). Comparing the Efficacy of Excitatory Transcranial Stimulation Methods Measuring Motor Evoked Potentials. Neural Plasticity, 2014, e837141. 10.1155/2014/837141

Moret, B., Donato, R., Nucci, M., Cona, G., & Campana, G. (2019). Transcranial random noise stimulation (tRNS): A wide range of frequencies is needed for increasing cortical excitability. Scientific Reports, 9(1), Article 1. 10.1038/s41598-019-51553-7

Moss, F., Ward, L. M., & Sannita, W. G. (2004). Stochastic resonance and sensory information processing: A tutorial and review of application. Clinical Neurophysiology, 115(2), 267–281. 10.1016/j.clinph.2003.09.014

Pavan, A., Ghin, F., Contillo, A., Milesi, C., Campana, G., & Mather, G. (2019). Modulatory mechanisms underlying high-frequency transcranial random noise stimulation (hf-tRNS): A combined stochastic resonance and equivalent noise approach. Brain Stimulation, 12(4), 967–977. 10.1016/j.brs.2019.02.018

Peirce, J., Gray, J. R., Simpson, S., MacAskill, M., Höchenberger, R., Sogo, H., Kastman, E., & Lindeløv, J. K. (2019). PsychoPy2: Experiments in behavior made easy. Behavior Research Methods, 51(1), 195–203. 10.3758/s13428-018-01193-y

Raveendran, R. N., Chow, A., Tsang, K., Chakraborty, A., & Thompson, B. (2021). Reduction of collinear inhibition in observers with central vision loss using anodal transcranial direct current stimulation: A case series. Brain Stimulation, 14(2), 207–208. 10.1016/j.brs.2020.12.015

Raveendran, R. N., Tsang, K., Tiwana, D., Chow, A., & Thompson, B. (2020). Anodal transcranial direct current stimulation reduces collinear lateral inhibition in normal peripheral vision. PLOS ONE, 15(5), e0232276. 10.1371/JOURNAL.PONE.0232276

Reinhart, R. M. G., Xiao, W., McClenahan, L. J., & Woodman, G. F. (2016). Electrical Stimulation of Visual Cortex Can Immediately Improve Spatial Vision. Current Biology, 26(14), 1867–1872. 10.1016/j.cub.2016.05.019

Rufener, K. S., Kauk, J., Ruhnau, P., Repplinger, S., Heil, P., & Zaehle, T. (2020). Inconsistent effects of stochastic resonance on human auditory processing. Scientific Reports, 10(1), Article 1. 10.1038/s41598-020-63332-w

Scolari, M., Kohnen, A., Barton, B., & Awh, E. (2007). Spatial attention, preview, and popout: Which factors influence critical spacing in crowded displays? Journal of Vision, 7(2), 7. 10.1167/7.2.7

Silva, A. E., Lehmann, R., Perikleous, N., & Thompson, B. (2023). The temporal dynamics of visual crowding in letter recognition: Modulating crowding with alternating flicker presentations. Journal of Vision, 23(10), 18. 10.1167/jov.23.10.18

Silva, A. E., Lyu, A., Leat, S. J., Khan, S., Labreche, T., Chan, J. C. H., Li, Q., Woo, G. C., Woo, S., Cheong, A. M. Y., & Thompson, B. (2022). A differential effect of visual cortex tDCS on reading of English and Chinese in patients with central vision loss. Brain Stimulation, 15(5), 1215–1217. 10.1016/j.brs.2022.08.016

Stagg, C. J., Best, J. G., Stephenson, M. C., O’Shea, J., Wylezinska, M., Kincses, Z. T., Morris, P. G., Matthews, P. M., & Johansen-Berg, H. (2009). Polarity-Sensitive Modulation of Cortical Neurotransmitters by Transcranial Stimulation. The Journal of Neuroscience, 29(16), 5202–5206. 10.1523/JNEUROSCI.4432-08.2009

Strasburger, H., Rentschler, I., & Jüttner, M. (2011). Peripheral vision and pattern recognition: A review. Journal of Vision, 11(5), 13. 10.1167/11.5.13

Terney, D., Chaieb, L., Moliadze, V., Antal, A., & Paulus, W. (2008). Increasing Human Brain Excitability by Transcranial High-Frequency Random Noise Stimulation. Journal of Neuroscience, 28(52), 14147–14155. 10.1523/JNEUROSCI.4248-08.2008

The jamovi project (2023). jamovi (Version 2.3) [Computer Software]. Retrieved from https://www.jamovi.org

van der Groen, O., Potok, W., Wenderoth, N., Edwards, G., Mattingley, J. B., & Edwards, D. (2022). Using noise for the better: The effects of transcranial random noise stimulation on the brain and behavior. Neuroscience & Biobehavioral Reviews, 138, 104702. 10.1016/j.neubiorev.2022.104702

Wallace, J. M., Chung, S. T. L., & Tjan, B. S. (2017). Object crowding in age-related macular degeneration. Journal of Vision, 17(1), 33–33. 10.1167/17.1.33

Whitney, D., & Levi, D. M. (2011). Visual crowding: A fundamental limit on conscious perception and object recognition. Trends in Cognitive Sciences, 15(4), 160–168. 10.1016/j.tics.2011.02.005

Wu, J., Yan, T., Zhang, Z., Jin, F., & Guo, Q. (2012). Retinotopic mapping of the peripheral visual field to human visual cortex by functional magnetic resonance imaging. Human Brain Mapping, 33(7), 1727–1740. 10.1002/hbm.21324

Yu, D., Cheung, S. H., Legge, G. E., & Chung, S. T. L. (2010). Reading speed in the peripheral visual field of older adults: Does it benefit from perceptual learning? Vision Research, 50(9), 860–869. 10.1016/j.visres.2010.02.006

